# A pipeline for cell migration analysis in live-cell imaging data from human iPSC-derived forebrain assembloids

**DOI:** 10.64898/2026.05.17.725711

**Authors:** Maya P. Weidman, Natalie Baker Campbell, Cody Headings, Samantha Chung, Musarat Khan, Aarnav Kandukuri, Vianne Lim, Gloria Olubowale, Michelle Kim, Anna Devor, Ella Zeldich, Martin Thunemann

**Affiliations:** Department of Biomedical Engineering, Boston University, Boston, MA, USA; Department of Anatomy & Neurobiology, Boston University Chobanian & Avedisian School of Medicine, Boston University, Boston, MA, USA; Neurophotonics Center, Boston University, Boston, MA, USA; Athinoula A. Martinos Center for Biomedical Imaging, Department of Radiology, Harvard Medical School, Massachusetts General Hospital, Charlestown, MA, USA; Center for Systems Neuroscience, Boston University, Boston, MA, USA

**Keywords:** forebrain assembloids, interneuron migration, oligodendrocyte migration, live-cell imaging, manual cell tracking, 4D image analysis

## Abstract

During forebrain development, inhibitory interneurons and oligodendrocyte progenitor cells migrate long distances into the developing dorsal cortex. Human induced pluripotent stem cell-derived forebrain assembloids (FAs) provide direct experimental access to this migratory process in vitro. Using viral labeling to express yellow fluorescent protein (EYFP) and tandem-dimer tomato (tdTomato) driven by EF1α or SOX10 promoters, respectively, we tracked cells in FAs over 15-17h using spinning disk confocal microscopy. We developed an end-to-end processing pipeline for 4D volumetric imaging data, consisting of background subtraction and drift correction, manual cell coordinate tracking, and an analysis workflow to describe migratory cell behavior. Image preprocessing significantly improved data quality for subsequent manual tracking in datasets with heterogeneous labeling density and brightness. Trajectory analysis of 336 EYFP- and 337 tdTomato-labeled cells from twelve FAs indicates that most cells show super-diffusive directed motility. Our pipeline represents a key resource for cell tracking in FAs and similar three-dimensional platforms. This pipeline represents the first open tracking resource for iPSC-derived FAs and can be used as a ground-truth resource for the development of automated cell detection and tracking algorithms.

## INTRODUCTION

The migration of neurons and glia from their birthplace to their final position in the developing telencephalon is one of the most spatially complex events in mammalian brain development. As such, gamma-aminobutyric (GABA)-ergic interneurons generated in the medial and lateral ganglionic eminence (MGE and LGE, respectively) must traverse long distances to populate the cerebral cortex, arriving between gestational weeks 8 and 15 in humans (Zecevic, Hu et al. 2011, Ma, Wang et al. 2013, Al-Jaberi, Lindsay et al. 2015). Oligodendrocyte progenitor cells (OPCs) originate from the same ventral domains and disperse broadly to myelinate axons across cortical and subcortical regions (Kessaris, Fogarty et al. 2006, Marton, Miura et al. 2019). Disruptions to these migratory programs arising from genetic perturbations, altered signaling environments, or intrinsic cellular deficits have been linked to cortical hypocellularity, imbalanced excitatory-to-inhibitory transmission, and deficient myelination in conditions such as Trisomy 21 (Down Syndrome), schizophrenia, autism spectrum disorders, Timothy Syndrome (Birey, Andersen et al. 2017), and epilepsy (Whittle, Sartori et al. 2007, Guidi, Ciani et al. 2011, Pan, Wu et al. 2019). Despite the clinical significance of these events, directly observing human cell migration in a tractable experimental system has remained challenging.

Assembloid models are generated by fusing regionally patterned organoids derived from human induced pluripotent stem cells (iPSCs) and provide a uniquely powerful model for studying interregional cell migration in vitro (Birey, Andersen et al. 2017, Pasca 2018, Sloan, Andersen et al. 2018). Forebrain assembloids generated by fusing ventrally patterned cortical organoids (vCOs) that contain mostly GABAergic inhibitory neurons and OPCs (Marton, Miura et al. 2019) with dorsally patterned cortical organoids (dCOs) that contain primarily glutamatergic excitatory neurons (Pasca, Sloan et al. 2015), recapitulate the pallial-subpallial boundary at the fusion interface. Upon physical assembly, virally labeled cells from the vCO region migrate into the dCO region, producing trajectories that parallel the ventral-to-dorsal migration program observed *in vivo* (Bagley, Reumann et al. 2017, Kim, Xu et al. 2019, Miura, Li et al. 2022), capturing the biological context of GE-derived cell migration within a controlled *in vitro* setting accessible through live-cell imaging. Selective fluorescent labeling of vCO-derived cells – most commonly achieved through lentiviral or adeno-associated virus (AAV)-mediated delivery of fluorescent reporter constructs under constitutive promoters such as EF1α or CAG for broad population labeling, or cell-type-specific promoters such as SOX10 or DLX1/2 for selective targeting of OPCs and GABAergic interneurons (Potter, Petryniak et al. 2009, Pol, Lang et al. 2013) – enables real-time visualization of migrating cells as they translocate from the vCO into the unlabeled dCO region during imaging (Bagley, Reumann et al. 2017, Birey, Andersen et al. 2017).

Four-dimensional (4D) spinning disk confocal microscopy (SDCM) enables the visualization of the cell migratory behavior over multi-hour imaging sessions, generating volumetric 4D datasets within intact assembloid volumes (Nakano 2002, Ewald, Werb et al. 2011, Oreopoulos, Berman et al. 2014). SDCM is particularly well-suited for these recordings because its parallel illumination architecture minimizes phototoxicity and photobleaching relative to single-beam scanning systems, enabling extended live imaging without compromising cell viability (Icha, Weber et al. 2017).

Despite these advantages, 4D assembloid imaging datasets present substantial analytical challenges not adequately addressed by existing tools. The complex 3D architecture of assembloid tissue introduces non-uniform background illumination gradients, light scattering and absorption with imaging depth, and pronounced differential fluorescence between the densely labeled vCO region and the sparsely labeled dCO region; multi-tile acquisition formats additionally introduce intensity discontinuities at tile boundaries. Progressive bulk drift of the assembloid relative to the imaging plane adds a further temporal artifact that is often detected in long-duration recordings. If left uncorrected, these factors obscure fluorescently labeled cell bodies and processes and introduce systematic error into downstream coordinate measurements. The densely packed, morphologically heterogeneous assembloid environment with cells ranging from compact spherical bodies to cells with elongated, filament-like processes poses severe challenges for existing automated segmentation and tracking algorithms, which are largely benchmarked on homogeneous datasets (Ulman, Maska et al. 2017, Maska, Ulman et al. 2023). Finally, no validated, publicly available pipeline or community reference implementation of migration metrics exists for 4D manual tracking data from brain assembloids; one of the field’s most comprehensive benchmarking resource, the Cell Tracking Challenge, contains no assembloid-derived 4D dataset (Ulman, Maska et al. 2017, Maska, Ulman et al. 2023). OrganoidTracker, the closest existing tool for organoid cell tracking, was trained on intestinal organoid and *C. elegans* embryo data and operates on 2D+time (single z-plane over time) rather than volumetric 4D data (Betjes, Kok et al. 2025), limiting its direct applicability to the dense, morphologically heterogeneous 4D assembloid imaging context addressed here.

Here, we present a validated, publicly available end-to-end framework designed to bridge these analytical gaps. The framework consists of three components: (1) a MATLAB pre-processing pipeline that converts raw SDCM data into background-subtracted BigTIFF hyperstacks; (2) a validated manual cell tracking workflow enabling precise 4D coordinate annotation; and (3) a MATLAB-based migration metrics code suite computing displacement, speed, directionality, and MSD-based motility descriptors from the resulting coordinate data. Applied to iPSC-derived FA data combining two fluorescent labeling strategies – a broad Ef1α-driven EYFP reporter and a SOX10 promoter-driven tdTomato reporter – the pipeline recovers biologically interpretable trajectories consistent with directed, superdiffusive migration across both labels. The coordinate dataset and code made available with this paper are intended as a practical resource for the community and a foundation for future automated tracking development in this system; access to the full imaging data, e.g., for machine learning training or algorithm benchmarking purposes is available upon reasonable request to the corresponding authors.

## MATERIALS AND METHODS

### iPSC lines and maintenance

Three iPSC lines were used: two female lines (WC-24-02-DS-B and ILD11-3), both thoroughly characterized in previous studies from our laboratory (Klein, Li et al. 2021, Li, Klein et al. 2022), and one male line (DS1-iPS4-disomic), generated by Dr. Orkin’s laboratory (Park, Arora et al. 2008, Maclean, Menne et al. 2012) and acquired through Boston Children’s Hospital. All lines exhibited normal karyotype. The lines were maintained on Matrigel (cat. 354277, Corning®, Corning, NY, USA) and passaged using ReLeSR (cat. 100-0483, STEMCELL Technologies, Vancouver, BC, Canada). mTeSR Plus medium (cat. 100-0276, STEMCELL Technologies) was replaced every other day. iPSCs at passages 24–50 with normal morphology were used for organoid generation.

### Cortical organoid generation and assembloid fusion

#### Dorsal cortical organoids (dCOs)

Oligodendrocyte-enriched dCOs were generated following the protocol of Madhavan, Nevin et al. (2018) with minor modifications. Briefly, iPSCs were dissociated with Accutase (cat. 7920, STEMCELL Technologies) and plated at 15,000 cells per well in low-adherence V-bottom 96-well plates (cat. MS-9096VZ, S-Bio Prime, Constantine, MI, USA) in 150 μL of mTeSR Plus supplemented with 50 μM ROCK inhibitor (cat. 12-541-0, Fisher Scientific, Pittsburgh, PA, USA). On day 1, medium was replaced with TeSR-E6 (cat. 0596, STEMCELL Technologies) supplemented with 2.5 μM Dorsomorphin (cat. P5499, MilliporeSigma, Burlington, MA, USA) and 10 μM SB-431542 (cat. S4317, MilliporeSigma), refreshed daily through day 6. From day 7, organoids were transitioned to Neural Medium (NM) containing Neurobasal A (cat. 10888022, Life Technologies, Carlsbad, CA, USA) with B-27 minus vitamin A (1:50; cat. 12587, Life Technologies), GlutaMax (1:100; cat. 35050061, Life Technologies), penicillin/streptomycin (1:100; cat. 15140122, Life Technologies), and Primocin (1:500; cat. ant-pm-1, Invivogen, San Diego, CA, USA). NM was supplemented with FGF2 (20 ng/mL; cat. 233-FB-25/CF, R&D Systems, Minneapolis, MN, USA) and EGF (20 ng/mL; cat. 236-EG-200, R&D Systems) between days 7–25. Cortical organoids were transferred to 24-well ultra-low attachment plates (cat. 07-200-602, ThermoFisher Scientific, Waltham, MA, USA) between days 16-24. BDNF (20 ng/mL; cat. AF-450-02, PeproTech, Waltham, MA, USA), NT3 (20 ng/mL; cat. 450-03, PeproTech), and 1% Geltrex (cat. A1569601, Life Technologies) were added from days 25-49. PDGF-AA (10 ng/mL; cat. 221-AA, R&D Systems) and IGF (10 ng/mL; cat. USA291-GF-200, R&D Systems) were added between days 50-60, followed by T3 (40 ng/mL; cat. T6397, MilliporeSigma) between days 61-69. From day 70, NM was changed every two days.

#### Ventral cortical organoids (vCOs)

vCO generation followed the dCO protocol with modifications to impose ventral forebrain identity (Marton, Miura et al. 2019). IWP-2 (5 μM; cat. S7085, Selleckchem, Houston, TX, USA) was added to TeSR-E6 between days 4–24. From day 7, vCOs were maintained in differentiation and maturation medium (DMM; Marton et al., 2019) composed of DMEM/F12 supplemented with B-27 minus vitamin A (1:50), N2 supplement (cat. 17502048, ThermoFisher Scientific), human insulin (25 μg/mL; cat. I9278-5ML, MilliporeSigma), NEAA (1:100; cat. 11140076, ThermoFisher Scientific), penicillin/streptomycin (1:100), GlutaMax (1:100), and β-mercaptoethanol (0.1 mM; cat. M3148, MilliporeSigma). EGF (20 ng/mL) and FGF2 (20 ng/mL) were added between days 7–24. SAG (1 μM; cat. 566660, MilliporeSigma) was added between days 12–24. Oligodendrocyte enrichment was achieved by supplementing DMM with BDNF (20 ng/mL), NT3 (20 ng/mL), PDGF-AA (10 ng/mL), IGF (10 ng/mL), T3 (60 ng/mL), HGF (5 ng/mL; cat. 315-23, PeproTech), cAMP (1 μM; cat. D0627, MilliporeSigma), and biotin (100 ng/mL; cat. B4639, MilliporeSigma) between days 24–37 (Marton et al., 2019). From day 37, vCOs were maintained in DMM supplemented with T3, cAMP, biotin, and ascorbic acid (AA; 20 μg/mL; cat. A4403, MilliporeSigma), with medium changes twice weekly.

#### Assembloid fusion

On day 75 of the differentiation, vCOs and dCOs were transferred into 1.5-mL microcentrifuge tubes containing 1 mL of medium and co-incubated for 4-8 days to allow physical adhesion. Assembloids were then transferred to 24-well ultra-low attachment plates and maintained with half-volume medium changes every other day (Birey, Andersen et al. 2017).

### Viral fluorescence labeling

Ten days prior to assembloid fusion (day 60 of differentiation), vCOs were labeled with AAV-DJ-Ef1α-eYFP (cat. GVVC-AAV-168, titer ≥1×10¹³ vg/mL, Stanford University Virus Core, Palo Alto, CA, USA), which drives EYFP expression under the ubiquitous EF1α promoter and pAAV-Sox10-tdTomato (titer ≥1×10¹³ vg/mL, cat. AAV1S(VB230712-1610aft)-K, VectorBuilder, Chicago, IL, USA) labeling GABAergic interneurons (Bagley, Reumann et al. 2017) and SOX10^+^ oligodendrocyte progenitor cells (Pol, Lang et al. 2013). Briefly, vCOs were transferred to 1.5 mL microcentrifuge tubes, medium was aspirated, and 0.5 μL of virus was added in 20-30 μL of medium for 30 minutes, followed by 300 μL of medium. vCOs were incubated overnight at 37°C and returned to 24-well plates until imaging.

### Immunohistochemistry

At day 82 of differentiation, selected vCOs were fixed for immunohistochemical validation of viral labeling efficiency. vCOs were fixed in 4% paraformaldehyde overnight, washed in PBS, and cryoprotected in 30% sucrose for 48 hours. vCOs were embedded in OCT compound (Fisher Scientific, cat. 23-730-571) mixed with 30% sucrose (60:40), flash-frozen, and cryosectioned into 12-μm sections. Sections were permeabilized and blocked in PBS containing 0.25% Triton X-100 (cat. 648466, MilliporeSigma) and 3% donkey serum (cat. D9663, MilliporeSigma) for 1 hour at room temperature (RT). Antigen retrieval was performed by microwaving slides twice at 25 °C for 5 minutes at 250W, allowing slides to cool to RT in between. Primary antibodies were incubated overnight at 4°C and secondary antibodies for 1 hour at room temperature, both in blocking solution. Slides were coverslipped with ProLong Gold Antifade Mountant with DAPI (cat. P36931, ThermoFisher Scientific). Primary antibodies: rabbit anti-GABA (1:250; cat. A2052, MilliporeSigma); goat anti-SOX10 (1:100; cat. AF3369, R&D Systems) used to characterize EYFP^+^ cell type identity. Secondary antibodies: anti-rabbit Alexa Fluor 750 (1:500; cat. ab175728, Abcam); anti-goat Alexa Fluor 633 (1:500; cat. A21082, ThermoFisher Scientific). Images were obtained using Slideview VS200 slide scanner (Evident Scientific, Waltham, MA, USA) with 20× objective and analyzed using QuPath v0.7.0 (Bankhead, Loughrey et al. 2017), counting cells labeled by each channel separately and measuring colocalization of channel markers. Representative tile scans were acquired on a Fluoview FV3000 confocal microscope with 40× oil immersion objective and deconvolved using cellSens imaging software. Stitching and background subtraction using Gaussian blur filtering was performed FIJI/ImageJ.

### Spinning disk confocal microscopy and image acquisition

Live 4D imaging of assembloids between day 110-120 of differentiation was performed on a Nikon CSU-W1 SoRA spinning disk confocal microscope equipped with a S PLAN FLUOR LWD 20× objective (0.7 NA, 2.3 mm working distance), controlled using NIS-Elements software. Imaging was conducted at 37°C with 5% CO₂ in a fully enclosed environmental chamber. EYFP-labeled cells were excited with a 488-nm solid-state diode laser and fluorescence was collected through a BP525/50 emission filter. tdTomato-labeled cells were excited with a 561-nm solid-state diode laser and fluorescence was collected through a BP609/54 emission filter. EYFP- and tdTomato-labeled cells were imaged within the same assembloid and up to 10 assembloids were imaged in parallel per recording session, using automated stage movement across wells of a 96-well glass-bottom plate.

For multi-tile acquisitions, adjacent fields were acquired with 15% overlap and stitched using NIS Elements built-in method. All remaining acquisition parameters are summarized in Table 1.

**Table 1.** Acquisition parameters for live-cell imaging.

| Parameter | Value |
| --- | --- |
| Microscope model | Nikon CSU-W1 SoRA spinning disk confocal |
| Objective / NA | S PLAN FLUOR LWD 20×, 0.7 NA, 2.3 mm WD |
| Lateral pixel size (XY) | 0.65 μm/pixel |
| Z-step | 20 μm |
| Number of Z-planes per timepoint | 16-19 |
| Time interval | 30 min |
| Total imaging duration | 17–18 hours |
| Laser excitation | 488 nm solid-state diode laser (EYFP), 561 nm solid-state diode laser (tdTomato) |
| Emission filter | BP525/50 (EYFP); BP609/54 (tdTomato) |
| Camera | Teledyne Photometrics Prime BSI Express 4.2MP Monochrome sCMOS |
| Bit depth | 16-bit |
| Raw file format | .ND2 (Nikon NIS-Elements software) |
| Imaging temperature | 37 °C (5% CO<sub>2</sub>) |
| Well format | 96 well glass bottom plate |

### Software and computational environment

All pre-processing and migration metric computations were performed in MATLAB (R2023b; The MathWorks Inc., Natick, MA, USA). Manual cell tracking and drift correction were conducted in FIJI/ImageJ v1.52p (Schindelin, Arganda-Carreras et al. 2012). Nikon ND2 files were read within MATLAB using the Bio-Formats library (Linkert, Rueden et al. 2010) via the bfGetReader interface. Gaussian filtering across timepoints was parallelized using the MATLAB Parallel Computing Toolbox. Computations were performed on a workstation equipped with an Intel Core i9-10920X CPU (3.50 GHz, 12 cores, 24 threads), 256 GB RAM, and a 10-TB SSD.

### Pre-processing pipeline

Raw 4D spinning disk confocal data underwent a four-stage pre-processing pipeline implemented in MATLAB prior to downstream tracking. Each stage is described below.

#### File ingestion and metadata extraction

Raw ND2 files were opened using the Bio-Formats MATLAB bfGetReader function (Linkert, Rueden et al. 2010). Global metadata – including series count, channel count, timepoint count, Z-plane count, and XY image dimensions – was extracted programmatically from the embedded metadata store. For data processing, one time series corresponding to a single assembloid field of view of multiple tiles within the multi-well acquisition in one channel color were selected prior to processing; all downstream steps operate on a single-channel, single-series dataset at a time. Raw image planes were read frame-by-frame using bfGetPlane and assembled into a 4-dimensional array with dimensions (X, Y, Z, T).

#### 3D Gaussian background subtraction

Long-duration live imaging of assembloids introduces spatially non-uniform background illumination gradients and substantial brightness heterogeneity between the densely labeled vCO- and the sparsely labeled dCO-region of the assembloid. To address this, a 3D Gaussian filter was applied to each timepoint volume to generate a smoothed, low-frequency background estimate, which was then subtracted:

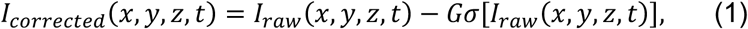

where Gσ denotes 3D Gaussian convolution applied with sigma values σ_x_ = σ_y_ = 21 pixels (27.3 μm) and σ_z_ = 9 pixels (180 μm), corresponding to FWHM values of FWHM_x_ = FWHM_y_ = 64.3 μm and FWHM_z_ = 424 μm. Gaussian filtering was applied in parallel across all timepoints using the MATLAB parfor function.

To quantify signal preservation, signal-to-background ratio (SBR) was computed using straight-line intensity profiles drawn in FIJI across fluorescently labeled structures surrounded by background. For each structure, the peak intensity along the profile was taken as the signal value and the histogram baseline flanking the peak was taken as the background value, giving SBR=peak/baseline for each measurement. SBR was measured across the same ten structures pre- and post-background subtraction per fluorescence channel and averaged to produce a single representative SBR value per condition.

#### Export to BigTIFF

Following pre-processing, 4D image stacks were exported as 16-bit BigTIFF hyperstacks using Bio-Formats bfsave with the BigTiff flag enabled. This format supports files exceeding 4 GB and maintains compatibility with FIJI/ImageJ and other software for downstream visualization and analysis. A singleton channel dimension was inserted by permuting the data array to (X, Y, Z, C, T) order prior to export.

#### Drift correction

Prolonged live imaging of freely floating assembloids produces progressive bulk drift of the sample relative to the imaging field across all three spatial dimensions, caused by stage motion, and other mechanical or thermal instabilities inherent to long-duration live imaging of freely floating tissue. Uncorrected drift introduces systematic errors into coordinate measurements in X, Y, and Z. Following export of MATLAB-processed stacks to BigTIFF, 3D drift correction was applied in FIJI/ImageJ using the Correct 3D Drift plugin (Parslow, Cardona et al. 2014). The plugin uses full-frame phase cross-correlation to compute cumulative XY and Z shift vectors at each timepoint relative to the first frame as a reference and applies these shifts to register the full 3D volume at every subsequent timepoint back to the reference. Full-frame cross-correlation is preferred over centroid-based approaches for assembloid data because it is insensitive to local shape changes in the assembloid over time and reflects net cumulative displacement from the reference rather than accumulated path length, avoiding inflation by back-and-forth jitter. The resulting drift-corrected 3D stacks served as the final input for all downstream manual cell tracking.

### Manual cell tracking workflow

#### OrthoTrack tracking

Manual tracking was performed using FIJI/ImageJ (v1.52p;(Schindelin, Arganda-Carreras et al. 2012) and OrthoTrack (Shvedov, Analoui et al. 2024). OrthoTrack extends FIJI’s native orthogonal view functionality to display synchronized XY, YZ, and XZ cross-sections updated in real time as cursor position changes, enabling 3D centroid localization. Originally developed and validated for studying adult neuron migration in *in vivo* zebra finch imaging data (Shvedov, Analoui et al. 2024), we demonstrate here that OrthoTrack transfers directly to 4D assembloid imaging data. The complete tracking workflow is as follows:

1. **Dataset loading.** Pre-processed BigTIFF hyperstacks were opened in FIJI. Brightness/contrast adjustments were made for display purposes only and did not alter pixel values.
2. **Cell identification and inclusion criteria.** Cells were eligible for tracking if they satisfied all the following criteria: (a) continuously identifiable across all timeframes of the recording window; (b) located at or near the vCO-dCO boundary region at the start of tracking; and (c) exhibiting net displacement directed away from the vCO region over the recording period.
3. **Orthogonal view navigation.** Orthogonal view mode was activated (Image > Stacks > Orthogonal Views), displaying XY, YZ, and XZ views with a shared crosshair. The annotator navigated through Z-planes and timepoints simultaneously across all three views to localize the 3D centroid of each target cell.
4. **ROI placement via OrthoTrack.** Once the cell centroid was identified, a point ROI was placed using the multipoint tool. OrthoTrack automatically assigned the ROI to the current timeframe, added it to the ROI Manager, and advanced to the next timepoint. This was repeated at every timeframe through the final timepoint.
5. **ROI set export.** Upon completion of a full trajectory, the ROI set was saved to disk in FIJI’s native format, with a separate file per cell or assembloid. Naming conventions within the ROI Manager to delineate individual cell trajectories within each saved file ensured that cell identity could be unambiguously recovered during coordinate extraction.
6. **Coordinate extraction.** FIJI’s Measure function was used for each ROI set to extract 3D spatial coordinates (X, Y, Z-slice) and timeframe index as a CSV file. One CSV file was generated per assembloid per annotator session and served as the direct input for MATLAB processing.

#### Multi-annotator compilation and unit conversion

Tracking was distributed across eight trained annotators. Individual CSV files were aggregated using the MATLAB, applying uniform unit conversions: lateral coordinates from pixels to micrometers (0.65 μm/pixel); Z-coordinates from slice number to micrometers (20 μm/slice, corresponding to the z step size); and timeframe indices to minutes [(frame − 1) × 30 min]. Compiled data were saved as Excel files organized by dataset, series, and channel. A total of 673 cells were tracked across 12 assembloids (WC-24-02-DS-B: n = 5; ILD11-3: n = 3; DS1-iPS4-disomic: n = 4) from 4 independent sessions yielding tracks of 336 EYFP^+^ cells and 337 tdTomato^+^ cells. Metrics were computed independently for each label.

### Migration metrics code suite

All migration metrics were computed from compiled OrthoTrack coordinate data in MATLAB. Individual cell trajectories were identified by t = 0 timestamps marking each new track within the combined results table. For visualization, all trajectories were origin-normalized (translated to [0, 0, 0] at t = 0) to facilitate direct comparison across cells originating at different assembloid positions.

#### Cumulative path length

Total path length was computed as the sum of 3D Euclidean distances between all consecutive coordinate pairs, where j indexes consecutive timeframe pairs, T is the total number of timeframes in the track, (x_j_, y_j_, z_j_) is the 3D position of the cell at frame j, and the square root term gives the Euclidean step distance between frames j and j+1.

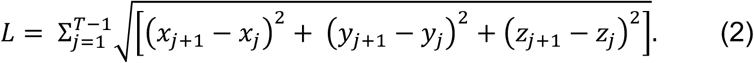

#### Net displacement

Net displacement was the straight-line Euclidean distance between the initial and final positions where (*x_1_*, y*_1_*, z*_1_*) and (x*_T_*, y*_T_*, z*_T_*) are the 3D positions of the cell at the first and final timeframes, respectively, and *D_net_* therefore reflects only the straight-line distance between start and end points regardless of the path taken in-between:

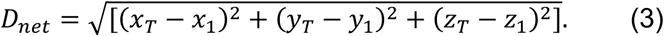

#### Instantaneous speed and speed variance

Frame-to-frame instantaneous speed was computed as 3D Euclidean distance between consecutive positions divided by the frame interval Δt = 30 min. For each cell, average frame-to-frame speed (̅v) and speed variance were computed over three temporal windows: the full imaging period, the first half, and the second half, providing a within-cell measure of temporal changes in motility not captured by global speed averages.

#### Directionality ratio

The directionality ratio (DR) was defined as:

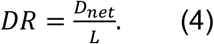

DR ranges from 0 (non-directional) to 1 (perfectly straight trajectory), providing a dimensionless measure of migration persistence independent of speed.

#### Mean squared displacement (MSD) and anomalous diffusion exponent

MSD analysis was performed using the *msdanalyzer* MATLAB toolbox (Tarantino, Tinevez et al. 2014). Tracks were organized as cell arrays of [time X Y Z] matrices and loaded using the msdanalyzer *addAll* interface. The time-averaged MSD was computed for each cell, where *r(t)* is the 3D position vector of the cell at time *t*, *τ* is the lag time*, |r(t+τ) − r(t)|^2^* is the squared 3D displacement between all timepoint pairs separated by *τ*, and <>*_t_* denotes averaging over all valid starting times *t* within the track:

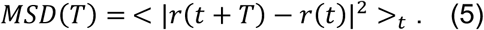

The population-level mean MSD curve was fit using a parabolic model consistent with directed diffusion:

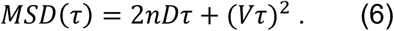

Where n=3 is the number of dimensions, D is the diffusion coefficient reflecting random motility, and V is the mean flow velocity reflecting the underlying directional drift component of migration. This model was selected over a simple linear fit based on the visibly parabolic shape of the population-level mean MSD curve, and its goodness of fit to that mean curve was confirmed by R^2^=0.998. To characterize per-cell migration mode, log-log fitting was additionally performed on MSD from individual cell curves using the first 25% of available lag times per track, fitting the power-law model:

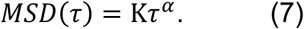

Where K is a generalized diffusion coefficient and α is the anomalous diffusion exponent. This approach restricts fitting to lag times where the MSD estimate is most reliable due to higher numbers of contributing track pairs. The resulting per-cell anomalous diffusion exponent α classifies individual cell migration mode: α ≈ 1 indicates Brownian (random diffusive) motion; 1<α<2 indicates superdiffusive (persistently directed) motion; and α ≈ 2 indicates ballistic motion (Dieterich, Klages et al. 2008).

### Code and coordinate data availability

To support adoption of the tracking framework and enable future benchmarking of automated tracking algorithms for brain assembloid data, we make the following resources publicly available alongside this paper: (1) all custom MATLAB pre-processing and migration metrics scripts; (2) the FIJI OrthoTrack workflow documentation; and (3) manually annotated cell coordinate data, provided both as raw FIJI tracking outputs organized by dataset and series, and as a single aggregated file with coordinates converted to physical units (µm, minutes) ready for direct use in trajectory analysis and metric computation. Full documentation of the repository structure and file formats is provided in the repository README. All code and coordinate data are deposited at https://github.com/codyheadings/ACMT.Raw image data and pre-processed hyperstacks are available upon reasonable request from the corresponding authors.

### Statistical analysis

All migration metrics were computed independently for each tracked cell and reported as pooled distributions within each channel: n=336 EYFP^+^ cells and n=337 tdTomato^+^ cells, each across 12 assembloids from 4 independent imaging sessions. Metrics are reported separately per channel and are not pooled across reporters. No inferential statistical comparisons between channels are performed in this paper; the dual-channel analysis is presented to demonstrate pipeline applicability and validate biological coherence of the tracking workflow across two distinct labeling strategies. Prior to computing summary statistics, outliers were identified and removed independently for each metric using the ROUT method (Q=1%; (Motulsky and Brown 2006)). Summary statistics are reported as median and the percentage of cells falling within defined histogram bins; median is preferred over mean given the right-skewed nature of most metric distributions, and bin-based percentages are reported to directly reflect the frequency distributions. All analyses were implemented in MATLAB (R2023b).

## RESULTS

### The assembloid model establishes the ventral-to-dorsal migration substrate

Forebrain assembloids were generated by fusing co-labeled EYFP^+^ and td-Tomato^+^ vCOs with unlabeled dCOs at day 75 of differentiation, following the regional patterning strategy illustrated in **Fig. 1**. In this system, EYFP^+^ cells and tdTomato^+^ cells are of ventral cortical origin and include GABAergic inhibitory interneurons and OPCs migrating from the vCO into the dCO upon fusion (**Fig. 1A–C**). SOX10 and GABA immunostaining confirmed the presence of both OPC-lineage cells and inhibitory GABAergic neurons within vCOs (**Supplemental Figure 1**). Our immunohistochemical analysis revealed that among EYFP^+^ cells, 51.47% of the cells were GABA positive, consistent with interneuron identity, while 46.78% were positive for SOX10, indicative of oligodendrocyte lineage. A fraction of EYFP^+^ cells (30.34%) co-expressed both GABA and SOX10 (**Supplemental Figure 1A, B**). Among tdTomato^+^ cells under the oligodendrocyte-specific SOX10 promoter, 49.29% of the cells were SOX10 positive, consistent with the oligodendrocyte identity (**Supplemental Figure 1A, C**). Live 4D SDCI revealed active translocation of EYFP^+^ and td-Tomato^+^ cells from the vCO region into the dCO region across the 17-18-hour recording window (**Fig. 1C-F**), confirming that the assembloid model produces the expected interregional migration substrate.

**Figure 1.**
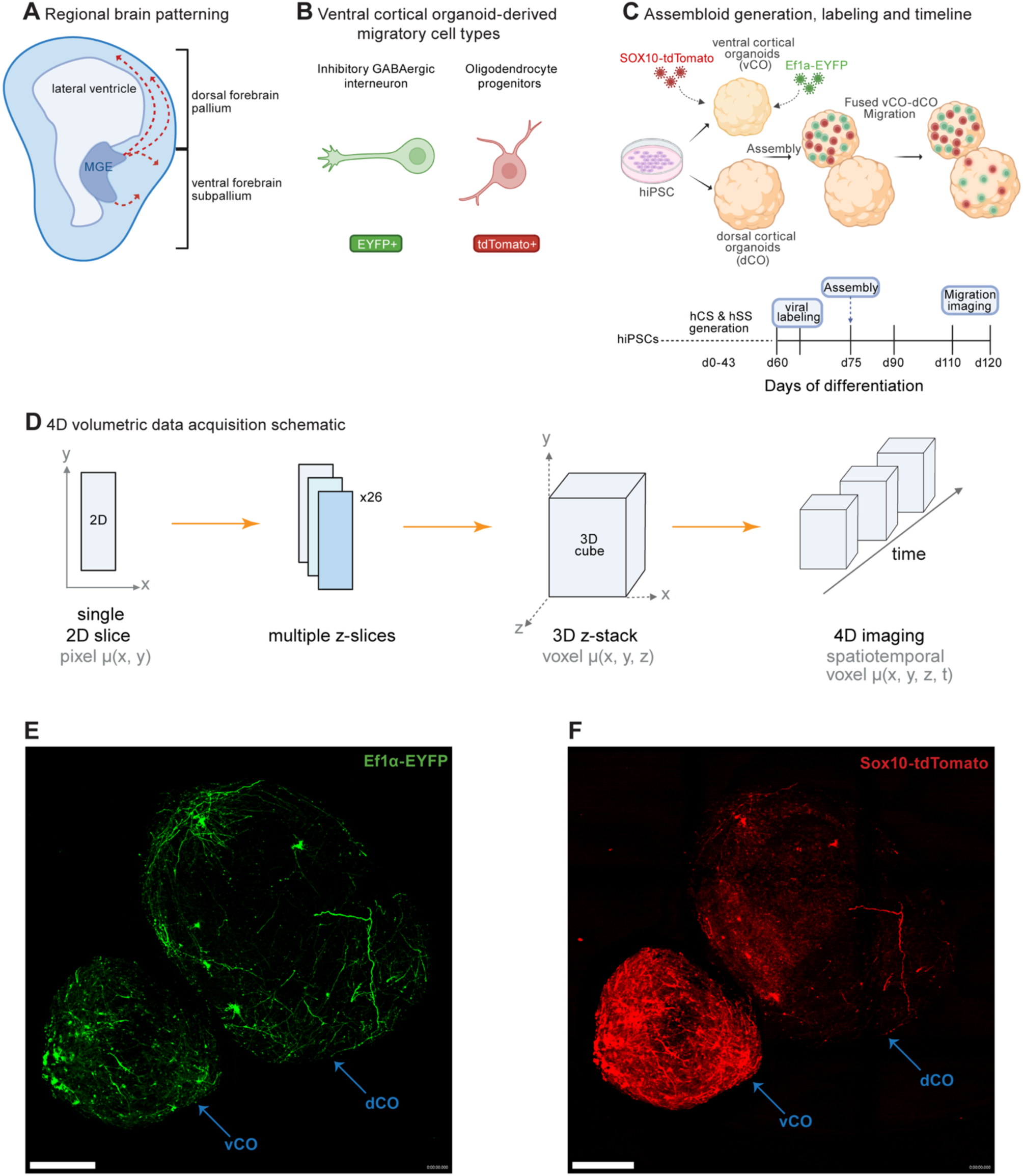
Human iPSC-derived forebrain assembloid model, cell labeling, and 4D imaging context. (A) Schematic of *in-vivo* interneuron and OPC migration from the medial ganglionic eminence (MGE) of the ventral subpallium to the dorsal pallium during human forebrain development. Red dashed arrows indicate migratory trajectories. (B) Ventral cortical organoid-derived migratory cell types modeled in this system: GABAergic inhibitory interneurons (EYFP^+^) and oligodendrocyte progenitor cells (tdTomato^+^). (C) Assembloid generation, viral labeling, and imaging timeline. vCOs were labeled with AAV-DJ-EF1α-eYFP (EF1α-EYFP) and AAV-SOX10-tdTomato (SOX10-tdTomato) on day 60 and fused with dCOs on day 75 to generate forebrain assembloids. (D) Schematic of 4D volumetric data acquisition: individual 2D z-slices (pixel μ(x,y)) are combined across 16 or 19 planes into a 3D z-stack (voxel μ(x,y,z)), one or more which are acquired at each timepoint to generate a 4D spatiotemporal dataset (voxel μ(x,y,z,t)). Z-stacks were acquired every 30 minutes for 17-18 hours. (E, F) Representative maximum intensity projections of a fused forebrain assembloid showing EF1α-EYFP^+^ cells (E, green) and SOX10-tdTomato^+^ cells (F, red). Blue arrows indicate the vCO and dCO regions of the assembloid. Scale bars: 400 μm.

### Pre-processing pipeline corrects background gradients and reduces bulk drift

Raw 4D data showed a pronounced brightness gradient between the densely labeled vCO and the sparsely labeled dCO portions of the assembloid (all virally-labeled cells present in the dorsal part have migrated from the ventral part), non-uniform background illumination across the field, and tile-boundary intensity discontinuities in multi-tile acquisitions (**Fig. 2A-C**, **Supplemental Fig. 2A-C**). Application of 3D Gaussian background subtraction equalized background intensities and improved the visibility of both EYFP- and tdTomato-labeled cells and processes in the dCO region (**Fig. 2C, Ci**, **Supplemental Fig. 2C, Ci**). The 3D Gaussian kernel has FWHM dimensions of 64.3 μm × 64.3 μm × 424 μm, which exceeds the size of individual cellular structures but captures low-frequency illumination gradients and the pronounced brightness imbalances between the densely labeled vCO and the sparsely labeled dCO region (**Fig. 2B, Bi, Supplementary Fig. 2B, Bi**). The resulting background-subtracted images isolate high-frequency cellular structures with sharp edges and well-defined boundaries, enhancing both visual interpretability and compatibility with downstream quantitative analysis and tracking workflows.

**Figure 2.**
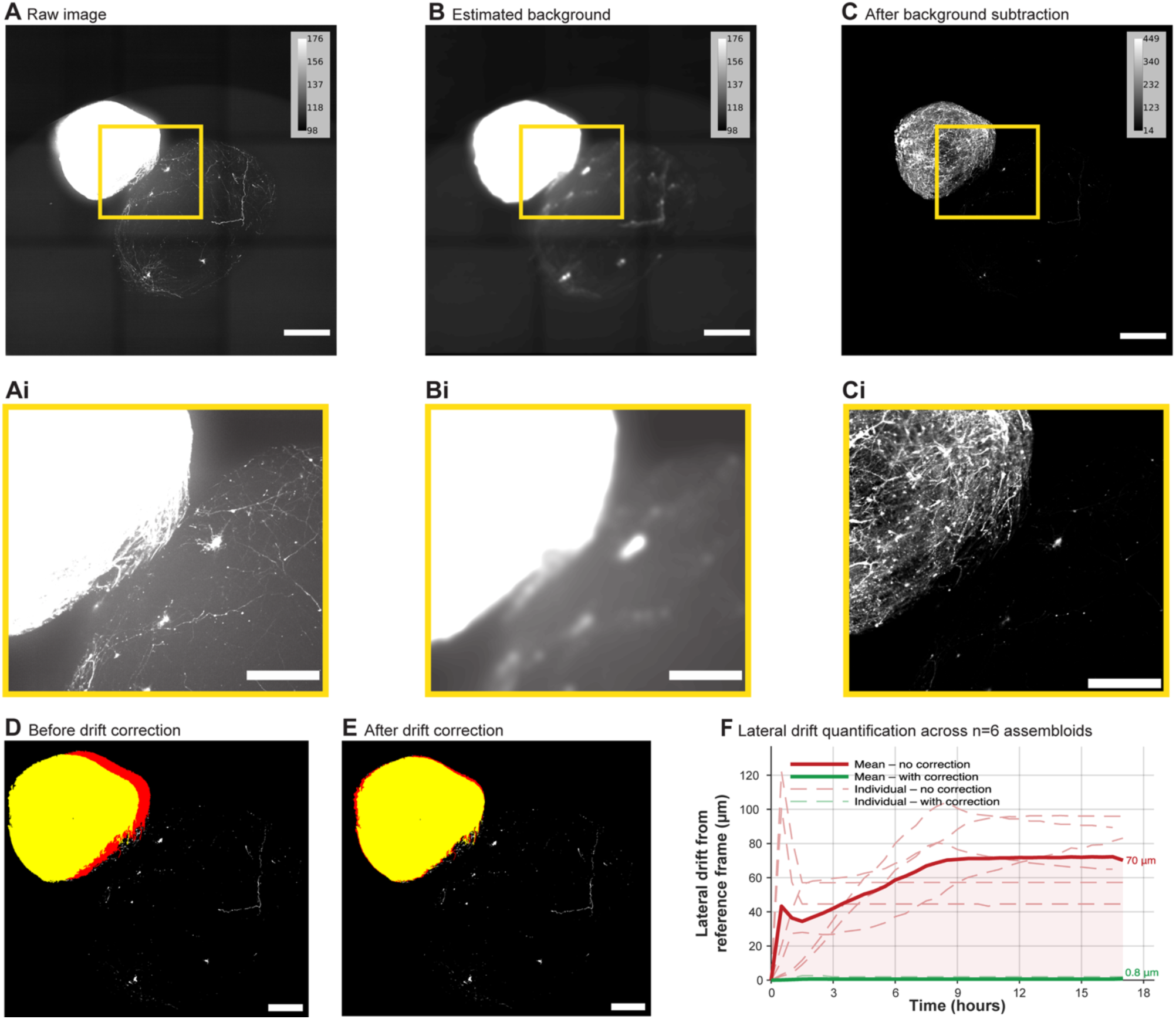
The pre-processing pipeline reduces background signal heterogeneity and corrects bulk lateral drift in 4D assembloid recordings. (A–C) Maximum intensity projections (z-MIP) of a representative assembloid (EYFP^+^ cells) at a single timepoint: raw image data (A), estimated background generated by 3D Gaussian filtering (FWHM: 64.3 × 64.3 × 424 µm) (B), and background-subtracted output (C). Intensity calibration bars are shown for each panel; panels A and B have matched windows (98-176) with narrow dynamic range; panel C is displayed over its native range (14-449). Yellow boxes indicate the region shown in Ai-Ci. (Ai-Ci) Zoomed view of the region indicated in A-C, displayed at identical brightness/contrast settings within the inset row. Equivalent results for the SOX10-tdTomato^+^ channel are shown in **Supplementary** Figure 2. (D, E) False-color overlay of the assembloid position at the first (yellow) and last timepoint (red) before (D) and after (E) drift correction, illustrating reduction in bulk lateral displacement. (F) Cumulative lateral drift magnitude from the reference frame as a function of elapsed time for n = 6 assembloids. Bold lines indicate group means; dashed lines indicate individual assembloid traces. Red: pre-processed but drift-uncorrected stacks; green: drift-corrected stacks. End-point mean values are annotated. Scale bars: 500 µm (A–C), 250 µm (Ai-Ci), 400 µm (D, E).

To quantify signal preservation, the signal-to-background ratio (SBR) was measured on ten fluorescently labeled structures in both the raw and background-subtracted images (**Supplementary Table 1**). For the EYFP channel, SBR improved approximately 39-fold from 5.4 in the raw data to 209.5 following background subtraction. In the same dataset, the SBR for the tdTomato channel improved approximately 47-fold from 1.9 in the raw data to 90.5 following background subtraction (**Supplementary Fig. 2**). These measurements confirm that 3D Gaussian background subtraction reduces background while preserving labeled cellular structure across two spectrally distinct fluorescent reporters.

To illustrate the magnitude of lateral bulk drift present in assembloid recordings of this duration, we applied the FIJI Correct 3D Drift plugin (Parslow et al., 2014) as a measurement tool to pre-processed stacks from six assembloids without drift correction, extracting cumulative XY shift vectors at each timepoint relative to the first frame. Lateral drift magnitude was computed as d(t) = √[dx(t)² + dy(t)²] × 0.65 µm/pixel. The mean cumulative lateral displacement reached 70 µm at the final timepoint with substantial variability between individual recordings, compared to an average lateral displacement of 0.8 µm in drift-corrected data (**Fig. 2F**). The considerable lateral bulk displacement in the absence of correction, motivated to implement drift correction as a standard component in the pre-processing pipeline.

### OrthoTrack workflow enables precise 4D manual cell tracking

Pre-processed and drift-corrected BigTIFF hyperstacks were loaded into FIJI/ImageJ and subjected to manual 4D cell tracking using OrthoTrack using the workflow outlined in **Fig. 3A**. The synchronized orthogonal view interface (**Fig. 3B**) enabled annotators to precisely localize fluorescently labeled cells in all three spatial dimensions simultaneously at every timeframe. The 3D coordinate assignment at each timepoint is conceptually illustrated in **Fig. 3C**, which depicts the anisotropic voxel geometry of the dataset (x, y, z dimensions: 0.65 µm x 0.65 µm x 20 µm) and the unique position recorded for the tracked cell at consecutive timepoints. The native output of OrthoTrack is a compressed FIJI ROI set from which 4D coordinate data are extracted by applying FIJI’s built-in Measure function to the loaded ROI set, generating a CSV file of X, Y, Z-slice, and timeframe indices in pixel units (**Fig. 3D**); these pixel-unit coordinates are subsequently converted to physical units by the MATLAB compile script.

**Figure 3.**
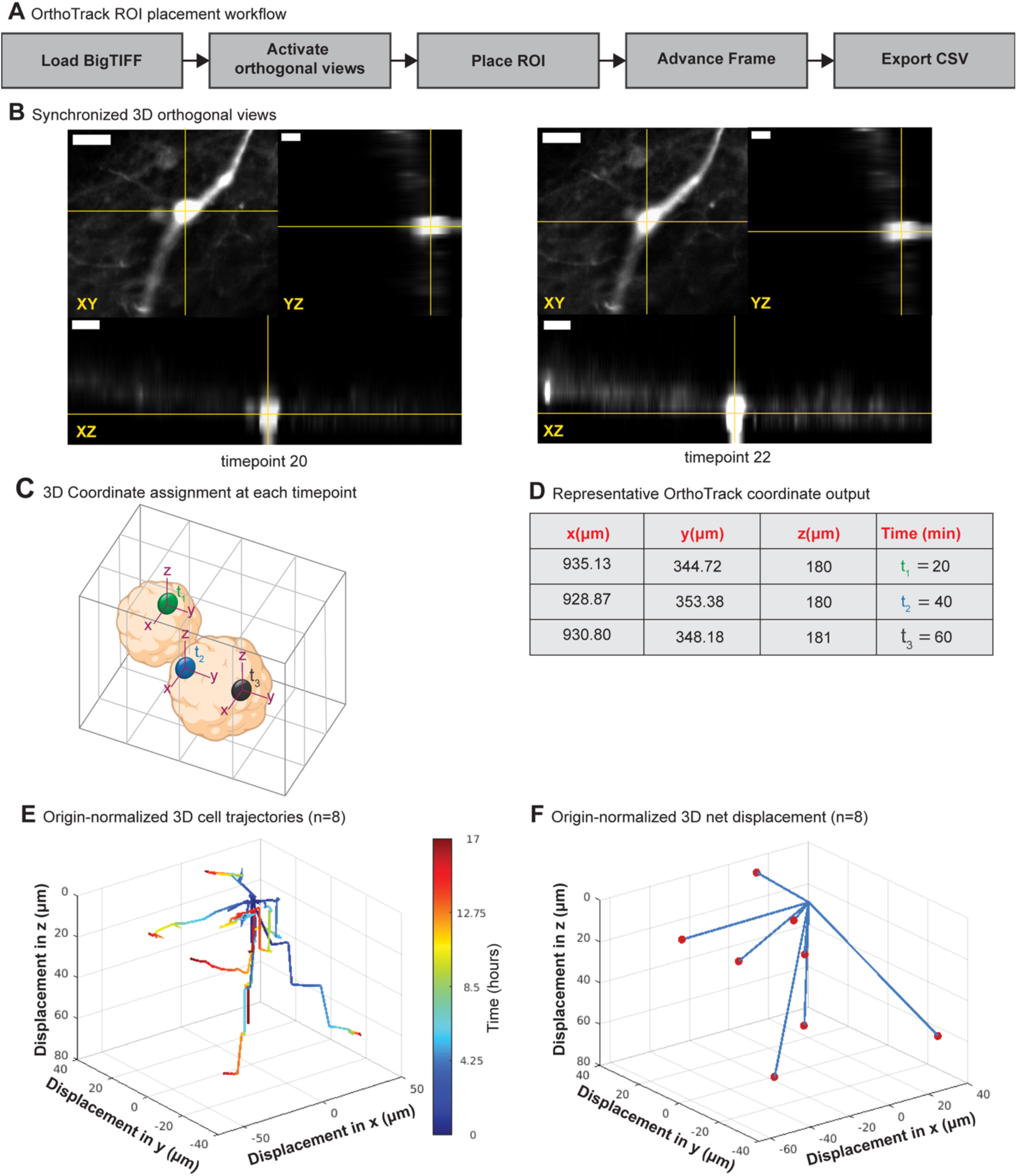
Manual 3D+time cell tracking workflow using OrthoTrack in FIJI/ImageJ. (A) Schematic overview of the OrthoTrack tracking workflow: pre-processed BigTIFF hyperstacks are loaded into FIJI/ImageJ, orthogonal views are activated to display synchronized XY, YZ, and XZ planes, point ROIs are placed at the cell centroid at each timeframe, the macro advances to the next frame, and the complete ROI set is exported as a CSV coordinate file. (B) Representative OrthoTrack interface screenshots showing synchronized XY, YZ, and XZ orthogonal views with yellow crosshair localization of a representative EYFP^+^ cell at two nearby timepoints (timepoint 20 and timepoint 22; 60-minute interval). Crosshair position is linked across all three views, enabling simultaneous 3D centroid localization. Scale bars: 20 µm (XY, YZ, XZ). (C) Schematic illustrating 3D coordinate assignment at each timepoint within the anisotropic voxel geometry of the dataset. The 3D bounding box reflects the coarser axial sampling (20 µm z-step) relative to the lateral pixel size (0.65 µm/pixel). A unique (x, y, z) coordinate is recorded for the tracked cell at t₁ (green sphere, vCO), t₂ (blue sphere), and t₃ (dark grey sphere, dCO), building a 3D+time trajectory across the assembloid volume. (D) Representative OrthoTrack coordinate output for a single tracked cell across three consecutive timepoints, showing 3D coordinates in physical units (µm) and elapsed time (minutes). (E) Origin-normalized 3D full trajectories of 8 representative tracked cells, color-coded by elapsed imaging time (hours). All trajectories are translated to a common origin at t = 0 to facilitate direct comparison of displacement patterns independent of absolute spatial position within the assembloid. (F) Origin-normalized 3D net displacement vectors for the same 8 representative cells shown in E. Blue lines connect the origin (t = 0) to each cell’s final position (red dot), illustrating endpoint displacement magnitude and direction. Together, panels E and F represent complementary visualizations of the 4D OrthoTrack coordinates which serve as the direct input to the migration metrics computed in Fig. 4. Axes in E and F indicate displacement in x, y, and z (µm) from the origin.

Origin-normalized 3D trajectory visualization of EYFP^+^ cells confirmed that tracked cells exhibit a diverse spectrum of motility-related behaviors: cells with large net displacements and relatively straight trajectories, as well as cells with more circuitous, exploratory paths, were both captured by the workflow (**Fig. 3E**). Net displacement, the vector connecting the cell’s original and final position further illustrate that this behavioral heterogeneity disperses across a range of directions and magnitudes within the assembloid volume (**Fig. 3F**). **Figure 3E-F** shows EYFP^+^ cells as a representative example of the 4D coordinate output. These coordinate outputs consisting of full trajectory paths and endpoint displacement vectors serve as the direct input to the quantitative migration metrics computed across both populations in **Fig. 4**.

**Figure 4.**
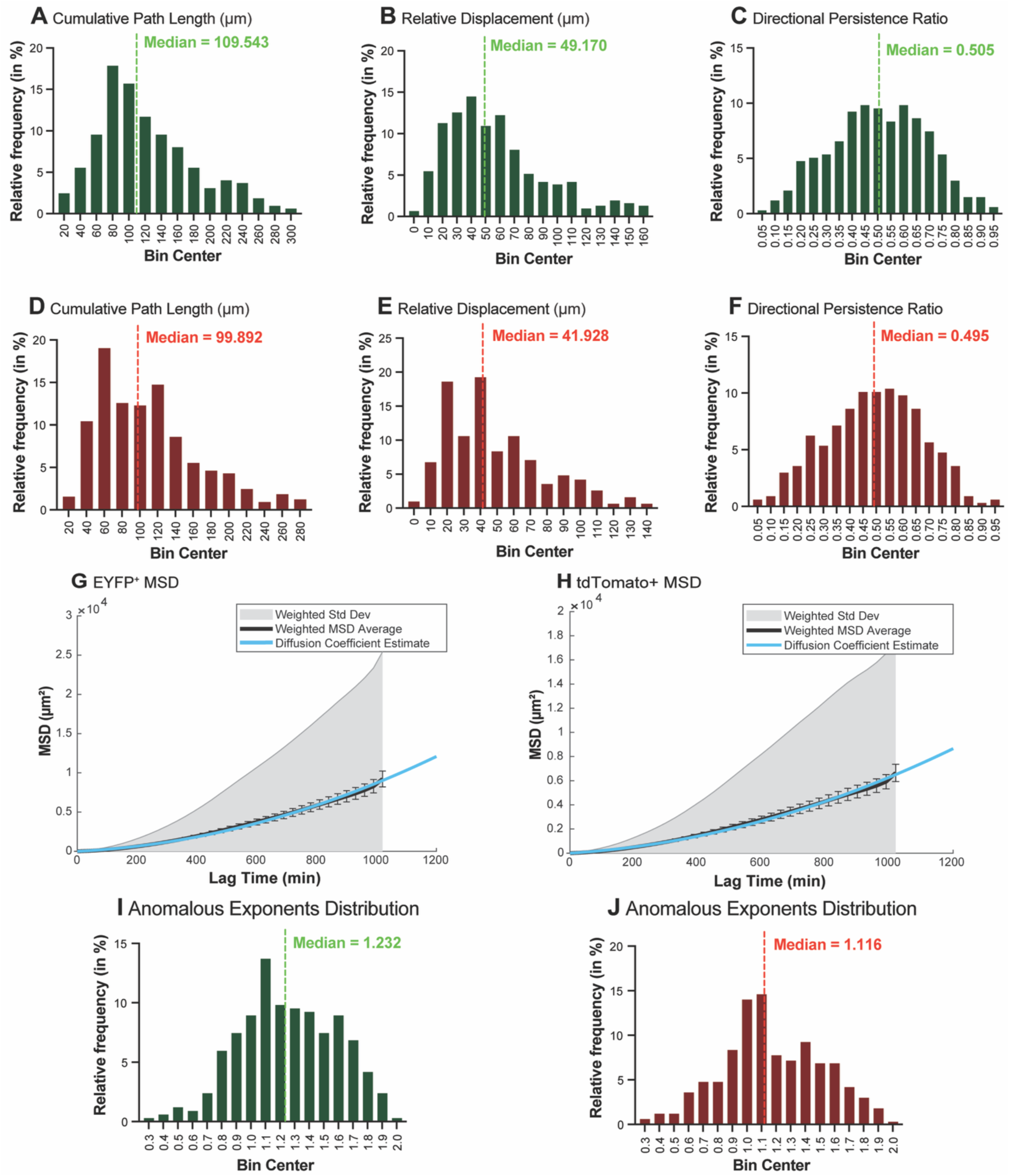
Migration metric distributions and MSD analysis across EYFP^+^ and tdTomato^+^ tracked cell populations. (A-C): n = 336 EYFP^+^ tracked cells. (D-F): n = 337 tdTomato^+^ tracked cells. All cells tracked across the same 12 assembloids from 4 independent imaging sessions. Distributions are shown as relative frequency histograms. Dashed lines indicate the median for each metric. Speed and speed variance distributions for both channels are shown in **Supplementary** Figure 3. (A, D) Cumulative path length (in µm): total 3D Euclidean path traveled by each cell across all consecutive timeframes. Right-skewed distributions with medians of 109.5 µm (EYFP^+^) and 99.9 µm (tdTomato^+^). (B, E) Net displacement (in µm): straight-line Euclidean distance between each cell’s initial and final 3D position. Right-skewed distributions with medians of 49.2 µm (EYFP^+^) and 41.9 µm (tdTomato^+^). (C, F) Directional persistence ratio (dimensionless, 0–1): ratio of net displacement to cumulative path length. Broad distributions across the full range with medians of 0.50 for both EYFP^+^ and tdTomato^+^. Values near 0 indicate circuitous, non-directional paths; values near 1 indicate straight, persistent trajectories. (G) MSD as a function of lag time (minutes) for 336 EYFP^+^ tracked cells. (H) MSD as a function of lag time for 336 tdTomato^+^ tracked cells. For both panels: gray shaded region: weighted standard deviation across all cell tracks; black line with error bars: weighted MSD mean; blue line: parabolic fit to the mean MSD curve. The upward-curving parabolic shape of the mean MSD curve is consistent with directed diffusion (R^2^=0.998 for both channels). EYFP^+^ fit: D=0.317µm²/min; tdTomato^+^ fit: D=0.276µm²/min.

### Migration metrics reveal biologically coherent, superdiffusive motility

Migration metrics for EF1α-EYFP-labeled cells are shown in **Fig. 4A-C** and for SOX10-tdTomato-labeled cells in **Fig. 4D-F**. MSD analysis is shown in **Fig. 4G, I** (EF1α-EYFP) and **Fig. 4H, J** (SOX10-tdTomato). Speed and speed variance distributions for both channels are provided in **Supplementary Fig. 3**.

Cumulative path lengths are broadly distributed with a right-skewed tail, with medians of 109.5 µm (EF1α-EYFP^+^; **Fig. 4A**) and 99.9 µm (SOX10-tdTomato^+^; **Fig. 4D)**. Approximately 78% of cells in both populations fell within 60-180 µm bins (77.8% for EF1α-EYFP^+^, 77.3% for SOX10-tdTomato^+^), with a sparse tail extending to approximately 300 µm. Net displacement showed similarly right-skewed distributions with medians of 49.2 µm (EF1α-EYFP^+^; **Fig. 4B**) and 41.9 µm (SOX10-tdTomato^+^; **Fig. 4E**); approximately 85% of EF1α-EYFP^+^ cells and 90% of SOX10-tdTomato^+^ cells displaced less than ∼90 µm from their origin over the 17-hour recording period. The consistently lower net displacement relative to cumulative path length across both populations reflects the degree of trajectory tortuosity.

The directionality ratio (DR) was broad and relatively uniformly distributed across the possible range, with a median of 0.50 for both cell populations (**Fig. 4C**, **F**). Migration solely by random diffusion would lead to a population clustered close to zero; here, a substantial proportion of cells maintain moderate-to-high directional persistence over the full recording window, alongside a minority of cells exhibiting more exploratory, low-persistence behavior.

Average instantaneous speed and its variance for both channels are provided in **Supplementary Fig. 3**. For the EF1α-EYFP population, speed showed a unimodal, approximately symmetric distribution centered near 3.65 µm/min (**Supplementary Fig. 3A**), while SOX10-tdTomato-labeled cells show a comparable distribution centered near 3.33 µm/min (**Supplementary Fig. 3C**). Variance of migration speed was markedly right-skewed in both channels, with medians of 0.021 (µm/min)^2^ for EF1α-EYFP (**Supplementary Fig. 3B**) and 0.018 (µm/min)^2^ for SOX10-tdTomato (**Supplementary Fig. 3D**). The relatively low speed variance observed across both populations is consistent with the 30-minute frame interval used in this study, which integrates over multiple saltatory migration cycles per timepoint; the burst-pause nucleokinesis characteristic of interneuron migration would require sub-minute temporal resolution to resolve at the individual step level (Tielens, Huysseune et al. 2016).

MSD analysis yielded population-level MSD curves with parabolic shapes for both cell populations, consistent with directed diffusion – a motility regime in which cells undergo both random diffusion and net directional drift simultaneously (**Fig. 4G, H**). For the EF1α-EYFP population (**Fig. 4G**), a parabolic fit to the mean MSD curve yielded an estimated diffusion coefficient D=0.317 µm²/min (95% CI: 0.285-0.349 µm²/min) and a mean flow velocity V=0.0825 µm/min (95% CI: 0.0807-0.0843 µm/min, R^2^=0.998), reflecting the net directional drift component of migration across the population. Per-cell log-log fitting of individual MSD curves confirmed superdiffusive behavior at the single-cell level (**Fig. 4I**): mean α = 1.253 ± 0.331 (s.d.) across 336 EYFP^+^ cells, with a median of 1.232 (Q1: 1.010, Q3:1.514). Most of these cells (76.5%) individually exhibited superdiffusive scaling (α>1), of which 26.2% showed strongly superdiffusive behavior (α>1.5); the remaining 23.5% of cells showed sub- or diffusive scaling (α≤1), reflecting the heterogeneity of the tracked EYFP^+^ population.

Prior to per-cell MSD fitting for the tdTomato^+^ population, one track was excluded on the basis that near-zero net displacement across the recording window produces insufficient signal for reliable log-log fitting; this cell was retained in all other metric distributions where near-stationary behavior is a valid and informative observation. MSD analysis therefore proceeded on 336 tdTomato^+^ tracks (**Fig. 4H**). Parabolic fitting yielded D = 0.276 µm²/min (95% CI: 0.251–0.302 µm²/min) and V = 0.0682 µm/min (95% CI: 0.0665–0.0698 µm/min; R² = 0.998). Per-cell log-log fitting yielded a mean α = 1.172 ± 0.342 (s.d.) across 336 tdTomato^+^ cells, median 1.116 (Q1: 0.952, Q3: 1.417), with 67.9% of cells individually superdiffusive (α > 1), of which 18.5% strongly superdiffusive (α > 1.5); 32.1% showed sub- or diffusive scaling (α ≤ 1) (**Fig 4J**). The consistency of superdiffusive MSD behavior across both fluorescent reporter populations confirms that the pipeline captures biologically meaningful directed motility regardless of the labeling strategy employed.

## DISCUSSION

We present a publicly available framework for quantitative volumetric cell migration analysis in iPSC-derived forebrain assembloids. Image pre-processing, manual tracking with OrthoTrack, and the migration metrics code suite each address complementary analytical gaps and together constitute a versatile, community-accessible resource for studying inter-region cell migration in brain assembloids, and related 3D live-imaging contexts. Rather than positioning automation as the immediate solution, we focus on establishing a robust and transparent framework for generating high-quality ground-truth data in challenging imaging conditions. To our knowledge, our pipeline generated the first 4D tracking dataset of two distinct virally labeled vCO-derived cell populations in the same assembloid.

### Pipeline development, implementation, and performance

The MATLAB pre-processing pipeline was a necessary step to enable accurate cell coordinate measurements by addressing challenges common to longitudinal live-cell imaging recordings of three-dimensional cultures. Specifically, we focused on two fundamentally distinct imaging artifacts: 3D Gaussian background subtraction corrects spatially structured illumination heterogeneity within individual timepoints, while full-frame cross-correlation-based drift correction removes bulk displacement across the recording window. The dimensions of the Gaussian kernel were chosen to be substantially larger than individual cell bodies across all three dimensions, preserving cellular structures while suppressing low-frequency background variation. Beyond improving visual quality for manual cell tracking, background subtraction improves data quality to be closer to the requirements of automated segmentation and tracking algorithms, which are particularly sensitive to background heterogeneity: detection thresholds calibrated for one region of a non-uniform field will systematically over- and under-detect cells in adjacent regions, producing false positives in bright areas and false negatives in dim ones. After background subtraction, we observed substantial SBR improvements for both fluorescent reporters, which serves as a validation metric for our current preprocessing steps, but also as a foundation necessary for automated approaches we and the field will eventually move toward.

We further observed that global drift magnitude varied substantially between recordings and was influenced by instrument configuration: assembloids were free-floating in culture media, and our microscope configuration moved the sample stage relative to a stationary imaging system. Configurations in which the objective moves while the sample remains stationary should reduce bulk motion of the assembloid; embedding assembloids in a matrix such as agarose prior to imaging is another option to restrict bulk motion. Where drift is severe, more robust volumetric motion correction algorithms such as NoRMCorre (Pnevmatikakis and Giovannucci 2017) may be preferable. We note that the quantification shown in **Fig. 2F** uses the same tool for both correction and measurement and should therefore be interpreted as demonstrating the scale of drift present in uncorrected data rather than as an independent performance validation.

The assembloid presents a challenging environment for cell tracking applications: high cell density, morphological heterogeneity spanning from compact spherical bodies to elongated filamentous structures, depth-dependent signal attenuation, and heterogeneous fluorescence. In our hands, pre-processed imaging data did not match the performance envelope of automated cell segmentation and tracking tools. Open-source and commercial tools using feature detection based on Difference-of-Gaussian, Laplacian-of-Gaussian, and ridge detection failed to produce reliable results on our datasets. Detectors based on intensity thresholding fragmented elongated cell processes into multiple false-positive objects, inflating counts and generating incoherent tracks, while tools operating on single z-planes did not produce the spatially and temporally linked 4D tracks required for migration analysis. Most importantly, in all cases, manual curation required to correct automated outputs exceeded the time cost of manual tracking, making manual annotation the more practical and reliable approach for this dataset. In line with this observation, the Cell Tracking Challenge documented that tracking accuracy on datasets with high fluorescence heterogeneity, irregular morphology, and dense packing remains substantially below performance on ‘cleaner’ data (Ulman, Maska et al. 2017, Maska, Ulman et al. 2023); our assembloid data likely falls into the challenging end of this spectrum.

The FIJI/ImageJ-based OrthoTrack plugin (Shvedov, Analoui et al. 2024) provides a meaningful ergonomic improvement over standard single-plane manual tracking approaches, and its application to human iPSC-derived assembloid models extends its validated use from tracking zebra finch neurons recorded in vivo with two-photon microscopy, demonstrating transferability to a different biological and optical context. The key feature of OrthoTrack in our hands was the synchronized three-view interface that constrains the annotator in all three dimensions simultaneously, providing sufficient coordinate accuracy independent of morphological complexity and the anisotropic nature of our datasets, where the voxel size in the axial direction (20 μm) is substantially larger than the lateral pixel size (0.65 μm). Tracking was distributed across eight independent annotators; formal inter-person reliability was not quantified, and post-hoc coordinate proximity analysis indicated that duplicate tracking of the same cell by different annotators occurred in approximately 7% of cases. A duplicate detection function with configurable proximity thresholds is provided in the deposited code repository and is recommended as a standard quality control step for future multi-annotator studies.

### Migration behavior of vCO-derived cells in forebrain assembloids

In the current study, we employed vCOs patterned to recapitulate MGE/LGE-like structures and enriched for OPCs (Marton, Miura et al. 2019). To label ventral lineage populations, we used a previously validated EYFP-reporter driven by the ubiquitous EF1α promoter (Birey and Pasca 2022), which labels a broad population of vCO-derived cells (Bagley, Reumann et al. 2017). Among EYFP⁺ cells, 67.92% expressed GABA and/or SOX10. Notably, immunohistochemical analysis revealed an unexpected overlap between GABA⁺ and SOX10⁺ populations (30.34%). This apparent co-expression may reflect several non-mutually exclusive biological scenarios: SOX10⁺ cells retain the molecular machinery required for GABA synthesis (Serrano-Regal, Bayon-Cordero et al. 2020), and such double-positive cells could represent transitional progenitor states arising from the ventral telencephalon. Consistent with this, OLIG2⁺ progenitors have been shown to give rise to GABAergic lineages, supporting the possibility of shared developmental intermediates marked by both oligodendrocyte- and interneuron-associated signatures (Ono, Takebayashi et al. 2008, Moyon and Casaccia 2017). In addition, 32.08% of EYFP⁺ cells did not co-localize with GABA or SOX10, suggesting labeling of additional populations under the EF1α promoter, potentially including astrocytes or excitatory neurons. Alternatively, these cells may represent immature interneurons that have initiated ventral patterning programs but have not yet acquired detectable GABA levels, potentially corresponding to DLX1/2⁺ progenitors (Potter, Petryniak et al. 2009, Sun, Pasca et al. 2016). We also observed only partial overlap between SOX10-tdTomato labeling and SOX10 immunostaining. This discrepancy may arise from differences between mRNA-driven reporter expression and protein-level detection, as well as potential limitations in antibody specificity and sensitivity. In addition, the fluorescent reporters used in this study label the entire cell volume, which contributed to the morphological heterogeneity that challenged both manual centroid localization and automated cell detection. In contrast, use of nuclear-localized fluorescent proteins (e.g., H2B-GFP) would confine the fluorescent signal to a compact, consistently spherical nucleus, producing more uniform object geometry for future automated segmentation while also reducing ambiguity in 3D centroid placement during manual tracking.

The migration metrics suite recovered from both labeled populations show similar behaviors. The directionality ratio distinguishes persistent migration from circuitous or exploratory behavior independent of speed. Sustained directional persistence, as indicated by the directionality ratio, is a key feature of interneuron and OPC migration at the population level: MGE-derived cortical interneurons maintain net directionality toward the cortex over extended periods, and ventrally derived OPCs co-migrate with interneurons along shared tangential routes in a directionally coordinated manner (Lepiemme, Stoufflet et al. 2022). Capturing this persistence metric at single-cell resolution is therefore not merely a technical validation output but reflects a biologically meaningful dimension of the migratory process. Reduced persistence of migrating cells can indicate impaired guidance, altered responsiveness to environmental cues, or disrupted organization of the cytoskeleton proteins. These motility-related features are observed in neurodevelopmental disorders, including Periventricular heterotopia (Fox and Walsh 1999), Lissencephaly (Squier and Jansen 2014), Polymicrogyria (Moon and Wynshaw-Boris 2013), as well as in inflammation-related processes (Boyd, Zhang et al. 2013, Smolders, Kessels et al. 2019). As such, persistence measurements provide a sensitive readout of migratory integrity that goes beyond simple measures of speed or displacement. The broad DR distributions reported here demonstrate the pipeline can faithfully recover this dimension of behavior across two distinct labeled populations. The population-level MSD curves for both channels follow parabolic rather than linear trajectories consistent with directed diffusion, while per-cell log-log fitting reveals that 76.5% of EF1α-EYFP^+^ and 67.9% and SOX10-tdTomato^+^ cells individually exhibit superdiffusive behavior (α > 1). Superdiffusive, directed-motion is consistent with the active migration of interneurons and OPCs under chemo-attractive guidance cues from the vCO toward the dCO (Dieterich, Klages et al. 2008, Shvedov, Analoui et al. 2024). In contrast, tracking noise would produce Brownian-like scaling (α ≈ 1) with no directional velocity component, while uncorrected bulk drift would produce near-ballistic scaling (α ≈ 2) uniformly across the population rather than the broad per-cell distribution observed here. The modest difference in α-value distribution between the two cell populations is consistent with the pre-eminent compositions of the two labeled populations – broadly labeled interneurons under the EF1α promoter versus OPCs under the SOX10 promoter. This result further underscores the value of simultaneous dual-channel tracking to capture the full spectrum of migratory behaviors present in the assembloid.

Most analyzed cells showed relatively low variance of speed; however, a minority exhibited more irregular, burst-like motion reflected in higher speed variance values. Importantly, data acquisition at a 30-minute interval was chosen in part to minimize phototoxicity over the 17-18-hour recording window. In contrast, individual saltatory migration cycles, which occur on minute timescales in migrating interneurons and OPCs (Tielens, Huysseune et al. 2016), were not resolved at this interval. The speed and speed variance metrics reported here therefore reflect migration rates averaged over the full recording window rather than instantaneous nucleokinesis dynamics.

The cell population represented in our tracking data is a selection-enriched subset of the total labeled population, as we annotated only actively migrating cells that were meeting all three inclusion criteria (continuous visibility across all frames, boundary-region location, and net directional displacement). Together with our observations on marker protein expression as discussed above, we therefore do not interpret our observations as definitive biological evidence of cell-type-specific migratory behaviors, lineage relationships, or cell state transitions. Rather, they highlight the complexity of marker-based classification in this system and establish a reproducible imaging and analysis framework for systematically interrogating ventral lineage populations. Future studies incorporating stage-specific reporters (e.g., DLX1/2-driven constructs) and orthogonal approaches will be required to more precisely resolve intermediate cell states (Birey and Pasca 2022).

In summary, the pipeline was applied without parameter modification to both EF1α-EYFP- and SOX10-tdTomato-labeled cells, demonstrating generalizability across labeling strategies and fluorophores. In preliminary tests, the pre-processing steps were directly applicable to data acquired with a different spinning disk confocal system (Yokogawa CQ1). We further anticipate that this pipeline will be adaptable to other types of assembloid and organoid models recapitulating cell motility, such as midbrain-striatum assembloids (Reumann, Krauditsch et al. 2023), tumor cell invasion datasets (Linkous, Balamatsias et al. 2019, Kim, Kim et al. 2025), as well as CRISPR-based screening studies targeting genes regulating cell migration and interactions with the extracellular matrix (Meng, Yao et al. 2023), which face equivalent background gradient and drift challenges. The manually generated ground-truth coordinate dataset can further serve as a gold standard for future development, optimization, and benchmarking of automated segmentation and tracking algorithms. As deep learning-based segmentation models trained on more biologically diverse datasets become available, automated approaches will become feasible – a transition the datasets and code released with this paper are intended to support.

## Declaration of generative AI and AI-assisted technologies in the writing process

During the preparation of this work the authors used Claude (Anthropic) to assist with language editing, proofreading, and improving clarity of the manuscript. After using this tool, the authors reviewed and edited the content as needed and take full responsibility for the content of the published article.

## ACKNOWLEDGEMENTS

The authors like to thank Naomi Shvedov (Ben Scott Laboratory and Neurophotonics Center, Boston University) for advice on OrthoTrack, the members of the Zeldich and Neurovascular Imaging Laboratories for helpful discussion. We thank Dr. Zahid Yaqoob and the members of the Boston University Neurophotonics Center for their help with the imaging setup. We thank the NIH for their support: NIH/NIA RF1AG088529 (MPIs: E. Zeldich and M. Thunemann), NIH/NINDS R21NS125469 (MPIs: E. Zeldich and M. Medalla), NIH/NIA and R03NS126864 (PI: E. Zeldich). We also thank A. Patil and S. Ehrhardt for their help with the manual tracking of cells in one dataset. We acknowledge support by the National Science Foundation (CBET 2215990). Research reported in this publication was supported by the National Science Foundation Award CBET 2215990. *The content is solely the responsibility of the authors and does not necessarily represent the official views of the National Institute of Health or the National Science Foundation*.

## AUTHOR CONTRIBUTIONS

**M. P. W.**, Methodology, Investigation, Software, Data curation, Formal Analysis, Visualization, Writing – original draft, Writing – review & editing; **N. B. C.**, Methodology, Investigation, Formal Analysis, Data curation, Visualization, Writing – review & editing; **C. H.**, Software, Formal Analysis, Data curation, Writing – review & editing; **S. C.**, **M. K.**, **A. K.**, **V. L.**, **G. O.**, **M. K.**, Formal Analysis; **A. D.**, Conceptualization, Supervision, Writing – review & editing; **E. Z.**, Conceptualization, Supervision, Funding acquisition, Investigation, Writing – original draft, Writing – review & editing; **M. T.**, Conceptualization, Software, Supervision, Funding acquisition, Writing – original draft, Writing – review & editing

## CONFLICT OF INTEREST STATEMENT

All authors declare that they have no conflicts of interest.

## Notes

https://github.com/codyheadings/ACMT

### Competing Interest Statement

The authors have declared no competing interest.

