## Supplementary Tables S1, Supplemental Figure 1,2,3 for "A pipeline for cell migration analysis in live-cell imaging data from human iPSC-derived forebrain assembloids"

### Data supplement

**Supplementary Table 1.** Estimation of signal-to-background (SBR) and signal-to-noise (SNR) ratios before and after background subtraction for EYFP- and tdTomato-labeled structures.

| EYFP | Before background subtraction | | | | | After background subtraction | | | | |
|---|---|---|---|---|---|---|---|---|---|---|
| **ROI** | **Signal** | **Back-ground** | **s.d. (back-ground)** | **SBR** | **SNR** | **Signal** | **Back-ground** | **s.d. (back-ground)** | **SBR** | **SNR** |
| 1 | 184.10 | 6.28 | 125.77 | 1.46 | 29.34 | 189.60 | 4.16 | 3.12 | 60.75 | 45.58 |
| 2 | 281.15 | 5.92 | 132.98 | 2.11 | 47.52 | 139.15 | 1.76 | 0.63 | 219.48 | 78.88 |
| 3 | 280.38 | 5.29 | 123.23 | 2.28 | 53.02 | 147.38 | 1.94 | 1.02 | 144.07 | 75.93 |
| 4 | 247.19 | 3.24 | 107.49 | 2.30 | 76.41 | 137.19 | 1.76 | 1.35 | 102.00 | 77.82 |
| 5 | 175.79 | 3.51 | 106.77 | 1.65 | 50.08 | 69.79 | 2.73 | 2.39 | 29.26 | 25.55 |
| 6 | 305.64 | 3.41 | 105.60 | 2.89 | 89.66 | 200.64 | 2.55 | 2.16 | 93.02 | 78.56 |
| 7 | 155.25 | 3.40 | 106.80 | 1.45 | 45.66 | 48.25 | 2.36 | 1.75 | 27.59 | 20.44 |
| 8 | 178.50 | 4.27 | 120.62 | 1.48 | 41.85 | 58.50 | 2.78 | 2.00 | 29.25 | 21.08 |
| 9 | 291.24 | 6.38 | 132.85 | 2.19 | 45.63 | 155.24 | 3.39 | 1.99 | 77.85 | 45.75 |
| 10 | 232.78 | 5.96 | 148.48 | 1.57 | 39.08 | 75.78 | 1.83 | 0.62 | 121.63 | 41.45 |
| **Mean** | | | | **1.94** | **51.83** | | | | **90.49** | **51.10** |
| | **Fold-improvement in SNR** | | | | | **0.99** | | | | |
| | **Fold-improvement in SBR** | | | | | **46.68** | | | | |

| tdTomato | Before background subtraction | | | | | Bfter background subtraction | | | | |
|---|---|---|---|---|---|---|---|---|---|---|
| **ROI** | **Signal** | **Back-ground** | **s.d. (back-ground)** | **SBR** | **SNR** | **Signal** | **Back-ground** | **s.d. (back-ground)** | **SBR** | **SNR** |
| 1 | 234.91 | 111.33 | 4.09 | 2.11 | 57.50 | 121.91 | 1.28 | 2.21 | 95.16 | 55.16 |
| 2 | 392.41 | 117.77 | 5.87 | 3.33 | 66.85 | 259.41 | 1.68 | 2.94 | 154.69 | 88.35 |
| 3 | 612.31 | 104.61 | 3.19 | 5.85 | 192.01 | 505.31 | 1.13 | 1.92 | 447.57 | 263.59 |
| 4 | 397.30 | 104.79 | 2.99 | 3.79 | 132.83 | 292.30 | 1.58 | 2.09 | 184.88 | 139.59 |
| 5 | 411.02 | 106.90 | 3.68 | 3.85 | 111.72 | 289.02 | 1.72 | 2.59 | 168.43 | 111.72 |
| 6 | 1467.58 | 247.37 | 22.51 | 5.93 | 65.19 | 1193.31 | 2.56 | 6.75 | 465.77 | 176.73 |
| 7 | 1575.71 | 279.36 | 38.23 | 5.64 | 41.22 | 1337.42 | 54.53 | 36.67 | 24.53 | 36.48 |
| 8 | 1988.03 | 239.52 | 39.83 | 8.30 | 49.91 | 1790.21 | 47.55 | 38.33 | 37.65 | 46.70 |
| 9 | 1773.87 | 160.18 | 14.26 | 11.07 | 124.44 | 1618.87 | 10.35 | 12.82 | 156.35 | 126.33 |
| 10 | 561.28 | 128.35 | 6.64 | 4.37 | 84.56 | 422.00 | 1.17 | 2.79 | 360.07 | 151.20 |
| **Mean** | | | | **5.43** | **92.62** | | | | **209.51** | **119.59** |
| | **Fold-improvement in SNR** | | | | | **1.29** | | | | |
| | **Fold-improvement in SBR** | | | | | **38.62** | | | | |

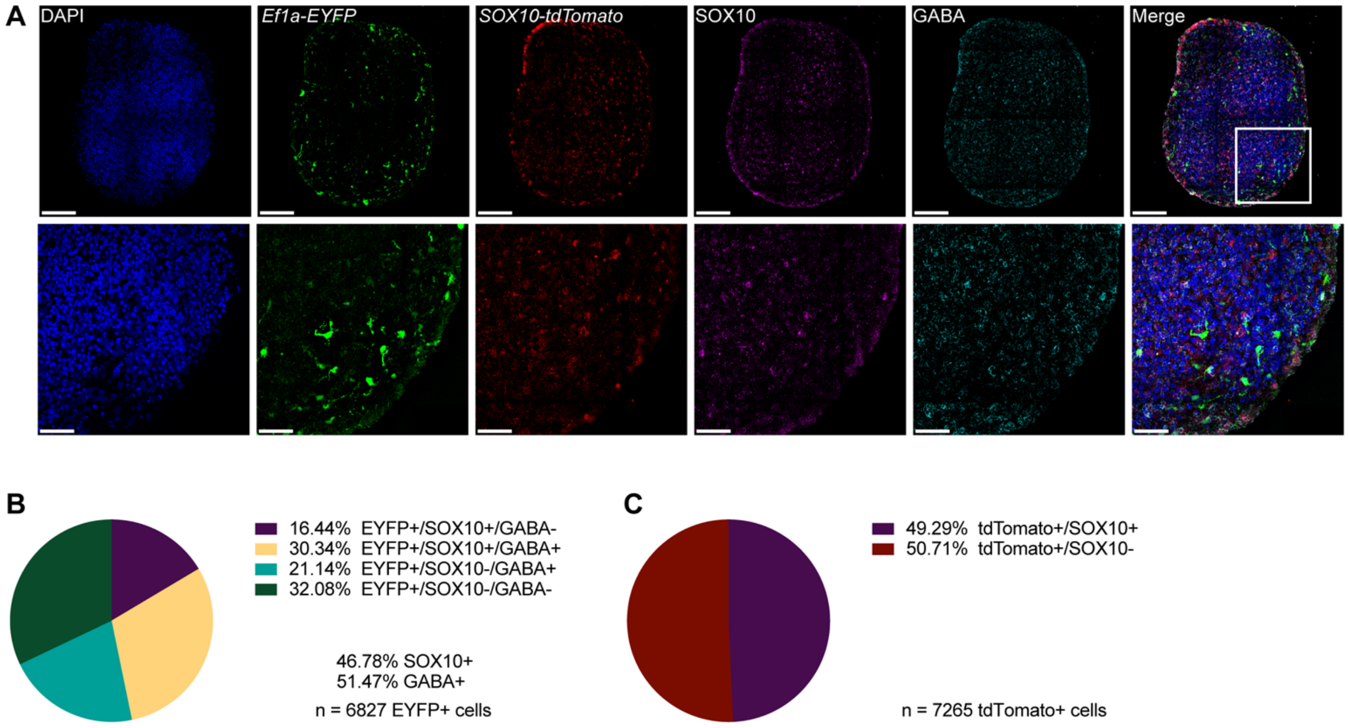


**Supplemental Figure 1: Efficiency of viral labeling in vCOs.** (A) Representative immunofluorescence images showing viral labeling of Ef1α-EYFP, SOX10-tdTomato, and immunohistochemical staining with SOX10, and GABA antibodies in vCOs at 82 days of differentiation. Scale bars are 150µm and 50µm (inset). (B, C) Quantification analysis was performed using QuPath across the same 3 vCOs. (B) Pie chart showing percentage breakdown of $EYFP^+$ cells colocalized with SOX10 and GABA. n = 6827 $EYFP^+$ cells. (C) Pie chart showing percentage breakdown of $tdTomato^+$ cells colocalized with SOX10. n = 7265 $tdTomato^+$ cells.

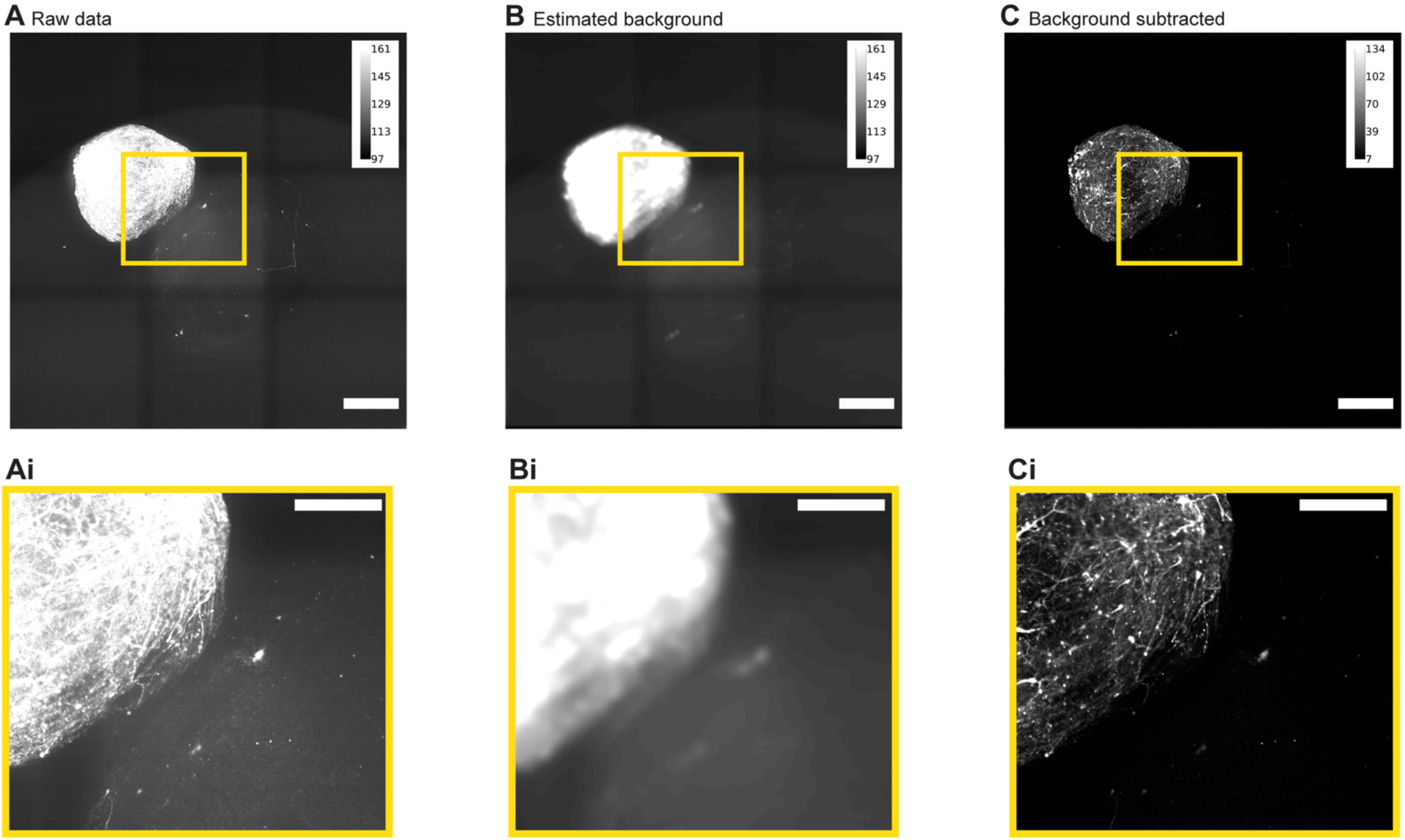


**Supplemental Figure 2: Background subtraction pipeline applied to Sox10-tdTomato$^{+}$ channel corrects signal heterogeneity.** (A–C) Maximum intensity projections (z-MIP) of the same representative assembloid shown in Figure 2 at a single timepoint, imaged in the red channel: raw data (A), estimated background generated by 3D Gaussian filtering (FWHM: 64.3 × 64.3 × 424 μm) (B), and background-subtracted output (C). Intensity calibration bars are shown for each panel. Yellow boxes indicate the region shown in Ai–Ci. (Ai–Ci) Zoomed insets of the indicated region, displayed at identical brightness/contrast settings within the inset row. Background subtraction markedly reduces illumination gradients and enhances visibility of SOX10-tdTomato$^{+}$ cell processes across the vCO and dCO regions. Signal-to-background ratio improved from 1.9 (raw) to 90.5 (corrected), approximately a 47-fold increase, measured from n = 10 SOX10-tdTomato$^{+}$ structures using straight-line intensity profiles drawn across the identical structures in both raw and background-subtracted images in FIJI/ImageJ. The same pipeline parameters were applied without modification from the EYFP$^{+}$ channel. Scale bars: 500 μm (A–C); 250 μm (Ai–Ci).

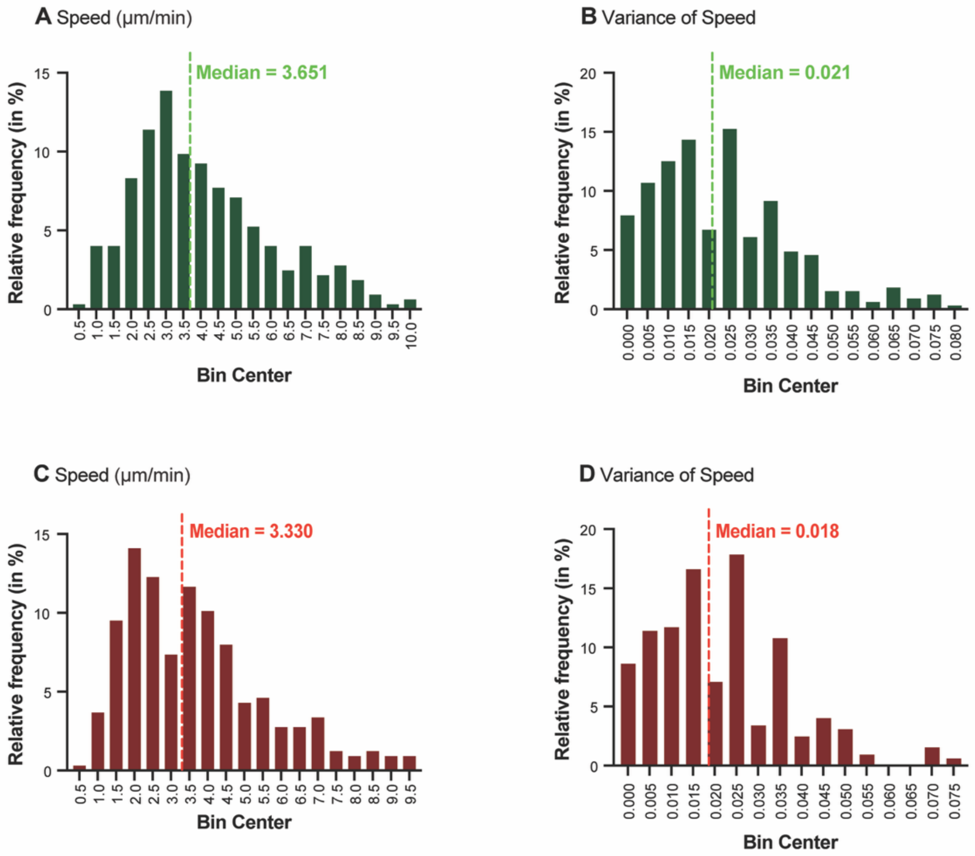


**Supplemental figure 3: Speed and speed variance histograms for both EYFP$^+$ and tdTomato$^+$ tracked cell populations.** (A, B): n = 336 EYFP$^+$ tracked cells. (C, D): n = 337 tdTomato$^+$ tracked cells. (A, C) Average instantaneous speed (μm/min): mean frame-to-frame speed computed as 3D Euclidean step distance divided by the 30-minute frame interval. Distributions are approximately unimodal with medians of 3.651 μm/min (EYFP$^+$) and 3.330 μm/min (tdTomato$^+$), with most cells falling between 1.5 and 6 μm/min in both channels. (B, D) Variance of speed ($(\mu m/min)^2$): variance of instantaneous frame-to-frame speeds across the full 17-hour imaging period. Markedly right-skewed distributions with medians of 0.021 $(\mu m/min)^2$ for EYFP$^+$ cells and 0.018 $(\mu m/min)^2$ tdTomato$^+$ cells, consistent with relatively steady migration paces in both populations.
